# The role of dermis resident macrophages and their interaction with neutrophils in the early establishment of *Leishmania major* infection transmitted by sand fly bite

**DOI:** 10.1101/2020.06.08.139956

**Authors:** Mariana M. Chaves, Sang Hun Lee, Olena Kamenyeva, Kashinath Ghosh, David Sacks

## Abstract

There is substantial experimental evidence to indicate that *Leishmania* infections that are transmitted naturally by the bites of infected sand flies differ in fundamental ways from the inflammatory and immune reactions initiated by needle inocula. We have used flow cytometry and intravital microscopy (IVM) to reveal the heterogeneity of sand fly transmission sites with respect to the subsets of phagocytes in the skin that harbor *L. major* within the first hours and days after infection. By flow cytometry analysis, dermis resident macrophages (TRMs) were on average the predominant infected cell type at 1 hr and 24 hr. By confocal IVM, the co-localization of *L. major* and neutrophils varied depending on the proximity of deposited parasites to the presumed site of vascular damage, defined by the highly localized swarming of neutrophils. Some of the dermal TRMs could be visualized acquiring their infections via transfer from or efferocytosis of parasitized neutrophils, providing direct evidence for the “Trojan Horse” model. The role of neutrophil engulfment by dermal TRMs and the involvement of the Tyro3/Axl/Mertk family of receptor tyrosine kinases in these interactions and in sustaining the anti-inflammatory program of dermal TRMs was supported by the effects observed in neutrophil depleted and in *Axl^-/-^Mertk^-/-^* mice. The *Axl^-/-^Mertk^-/-^* mice also displayed reduced parasite burdens but more severe pathology following *L. major* infection transmitted by sand fly bite.

**Summary:** Sand flies transmit *Leishmania major* which causes cutaneous leishmaniasis in humans and in non-human hosts. Our analyses of sand fly transmission sites of *L. major* in the mouse skin revealed that dermis resident macrophages (TRM) were the predominant phagocytes to take up the parasite within the first 24 hr post-bite. The early involvement of neutrophils varied depending on the proximity of deposited parasites to the site of tissue damage around which the neutrophils coalesced. By intra-vital microscopy, some of the dermal TRMs could be visualized acquiring their infections by direct transfer from or phagocytosis of parasitized neutrophils. The involvement of the Tyro3/Axl/Mertk family of receptor tyrosine kinases in these cellular interactions and in sustaining the anti-inflammatory functions of dermal TRMs was supported by the reduced parasite burdens but more severe pathology observed in *Axl^-/-^Mertk^-/-^* mice. The heterogeneity of sand fly transmission sites with respect to the dose of parasites and the early cellular interactions described here likely contribute to the wide range of infection outcomes that are associated with natural transmission of *L. major* observed in mouse models and possibly humans.

## Introduction

Kinetoplastid parasites of the genus *Leishmania* are phagosomal pathogens transmitted by phlebotomine sand flies that produce a spectrum of diseases in their human hosts, ranging from localized cutaneous lesions, to tissue destructive mucosal involvement, to disseminated, visceral disease. As would be expected for acquired resistance against pathogens that reside in a phagosome [1], there is ample evidence from experimental and clinical studies that T helper 1 (Th1) responses are a crucial component of the protective response. By contrast, there is less consensus regarding the types of phagocytes that harbor *Leishmania* in different tissues and in different stages of disease. The widely employed mouse models of cutaneous leishmaniasis, for example, have variously implicated neutrophils, inflammatory monocytes, monocyte derived dendritic cells (DCs) / macrophages, migratory dermal DCs, and dermis resident macrophages as cells that support transient or productive infections, or that kill the parasite in their constitutive or immune activated state [2-8]. These observations have frequently relied on infections initiated by relatively high dose inocula (10^5^-10^7^) delivered by needle in the skin or subcutaneous site.

There is now substantial evidence that infections initiated by needle differ in fundamental ways from those delivered by infected sand flies. Based on studies involving experimentally infected, laboratory-colonized sand flies, infection is typically established by fewer than 100 promastigotes, with occasional transmissions of a few thousand parasites by individual flies [9-11]. Sand flies can also co-egest other factors that are absent from needle inocula and that can modulate the host response, including saliva, microbiota, and components released by the parasites themselves, such as exosomes and promastigote secretory gel [12-15]. Infected sand fly bite sites are associated with substantial tissue damage and highly localized, inflammatory cell recruitment, characterized most prominently by neutrophils [13, 16]. *In vivo* imaging of sand fly transmission of RFP-*L. major* revealed that many of the deposited parasites were phagocytosed by neutrophils [16]. Furthermore, depletion of neutrophils prior to sand fly transmission was found to accelerate clearance of *L. major* from the site, suggesting that the acute neutrophilic, wound healing response to sand fly bite is exploited by the parasite to promote the infectious process [16]. This remains the only study to date to follow the fate of promastigotes following their natural delivery by sand fly bite. Because the studies were confined to two-photon intravital microscopy (2P-IVM) of the transmission sites using LYS-GFP mice, which favored direct visualization of neutrophils, the infection of other cell types and their interaction with neutrophils could not be easily addressed. In addition, these studies were undertaken prior to our ability to specifically label and image dermis resident macrophages, which based on recent observations regarding their high phagocytic capability and perivascular distribution in the steady-state dermis [3, 17], would seem to have a strong potential to engage transmitted parasites.

In the current studies, we have used flow cytometry and intravital imaging to reveal the heterogeneity of sand fly transmission sites with respect to the subsets of phagocytes that harbor *L. major* within the first hours and days after infection. We identified dermis resident macrophages as the predominant early population of infected cells that in some cases acquired their infections directly from parasitized neutrophils. The role of receptor tyrosine kinases in the interaction between neutrophils and dermis resident macrophages and the maintenance of their alternative activation phenotype during infection was also explored.

## Results

### Dermis-resident macrophages are the predominant infected population following transmission of *L. major* by sand fly bite

Our prior attempts to use flow cytometry to provide a more complete assessment of the subsets of infected cells in the transmission sites were unsuccessful due to the very low numbers of the RFP *L. major* Friedlin strain promastigotes deposited by the infected flies [10]. To address this question, we infected *P. duboscqi* sand flies with RFP *L. major* Ryan (RFP *LmRyn*) strain, which has been shown to produce infections in *P. duboscqi* that are associated with a far greater efficiency of successful transmissions by bite [18]. Flies infected with RFP *LmRyn* for 10-19 day were allowed to feed on the ears of C57Bl/6 mice which were processed for flow cytometric analysis of recovered cells at 1 hr, 24 hr, 5 d and 12 d post-bite (Fig 1A). As the ears were exposed to 10 infected flies for 2 hr in the dark, the 1 hr time point will reflect sites that were bitten 1-3 hr previously. RFP^+^ cells were detectable in the exposed ears of each of the 32 different transmission sites studied, with an average of 21 ± 10.4 and 49 ± 29.7 infected cells/10^5^ cells detected at 1 hr and 24 hr, respectively (Fig 1B). Based on our previously described gating strategy to identify the various RFP^+^ infected, myeloid populations in the skin (Fig 1C) [3], we observed that neutrophils, dermal tissue-resident macrophages (TRMs), inflammatory monocytes and monocytes-derived dendritic cells (mo-DCs) each became infected over the first 24 hours following transmission by bite (Fig 1D-F). Little if any role could be attributed to conventional DCs (cDCs) in the early infection by RFP *LmRyn*. Most striking was the wide range in the frequencies of infected cells that were either dermal TRMs (5-70%) or neutrophils (0-50%) in the different bite sites at these early time points (Fig 1E and F). On average, the dermal TRMs were the predominant population of infected cells at both 1 hr (38.1 ± 8.0%) and 24 hrs (46.0 ± 7.4%), and they were the only population to show a significant increase in the absolute numbers of infected cells during this time, from 9.6 ± 2.7 to 27.6 ± 8.0 per 10^5^ total cells. We attribute the increase in the absolute number of infected cells between 1hr and 24 hr as likely due to the transmitted parasites that had not yet been phagocytosed at 1 hr and that would have been excluded from the flow analysis of infected cells.

**Fig 1.**
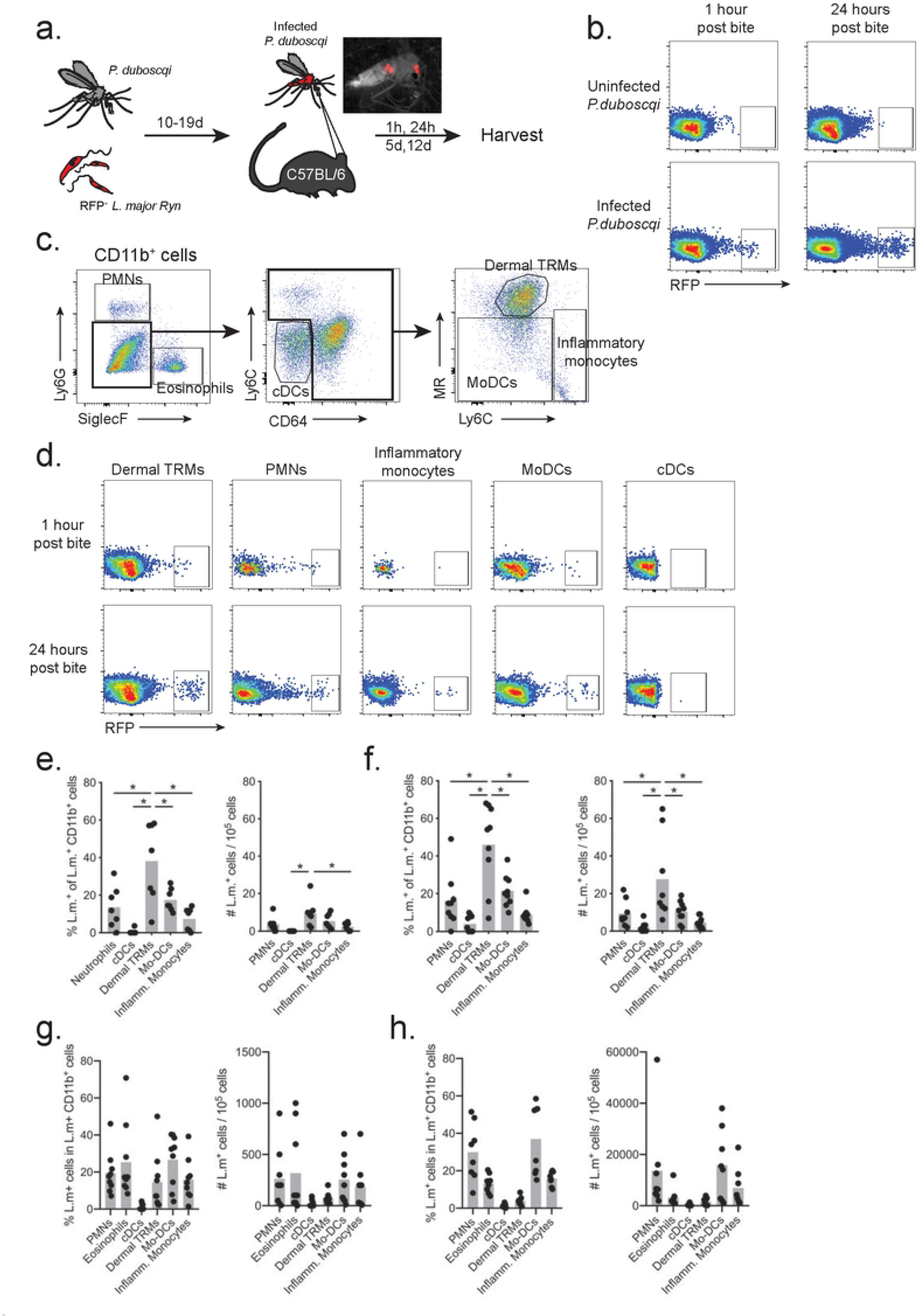
Infected skin phagocytes after sand fly transmission. (A) *Phlebotomus duboscqi* sand flies were infected with 5 x 10^6^ RFP^+^ procyclic promastigotes of *L. major* (*LmRyn*). After 10-19 days, C57Bl/6 mice ears were exposed to infected sand flies. Mice exposed to uninfected flies were used as controls. (B) Representative flow cytometric profiles obtained 1 hr and 24 hrs after sand fly exposure showing RFP^+^ and RFP^-^ CD11b^+^ cells recovered from the ear dermis. (C) Gating strategy performed on CD11b^+^ cells from naïve ear that was used for subset analysis of CD11b^+^RFP^+^ cells recovered from ears of mice exposed to infected sand flies: Ly6G^+^ (neutrophils), SiglecF^+^ (eosinophils), Ly6C^-^CD64^-^ (cDCs), Ly6C^-/inter^CD64^+^CD206^+^ (dermal TRMs), Ly6C^-/inter^CD64^+^CD206^-^ (mo-DCs) and Ly6C^+^CD64^+^CD206^-^ (Inflammatory monocytes). (D) Representative flow cytometric profiles of infected skin phagocytes 1 and 24 h post transmission. Frequency of infected cells and number of infected cells / 10^5^ CD11b^+^ cells at (E)1 hr, (F) 24 hr, (G) 5 days and (H) 12 days post-transmission. Values shown are percentages or number of cells in individual ears (closed circles), and mean percentages or number of cells per ear (grey bars), 6-8 ears per group pooled from 3 independent experiments; *\*p* < 0.05.

Five days following sand fly transmission, the total number of RFP^+^ CD11b^+^ cells had increased to 1113 ± 799 / 10^5^ cells, of which the frequency of infected cells that were dermal TRMs was reduced (14.6 ± 5.1%) in favor of neutrophils (19.4 ± 3.9%), mo-DCs (26.1 ± 4.7%), and inflammatory monocytes (15.9 ± 3.8%) (Fig 1G). Unexpectedly, 25.3 ± 6.8% of the RFP^+^ cells were eosinophils (Fig 1G). At 12 days post-transmission, the total number of RFP^+^ CD11b^+^ cells had increased to an average of 41,379 ± 38,843 / 10^5^ cells, of which neutrophils (30 ± 5.4%), mo-DCs (37.0 ± 6.6%), and inflammatory monocytes (15.9 ± 1.3%) remained the predominant infected cell types in the site, while there was a decline in the frequency of infected cells that were eosinophils (13.1 ± 1.8%) and dermal TRMs (3.8 ± 0.8%) (Fig 1H). Taken together, these data support an important role for dermal TRMs in the initial establishment of sand fly transmitted infections, with substantial variability observed in both the absolute number and relative frequencies of infected TRMs and neutrophils in different bite sites. By 1-2 weeks post-transmission, the infections transitioned to predominantly inflammatory cells, including neutrophils, eosinophils, monocytes, and monocyte-derived cells.

### Intravital microscopy of dermal TRMs and neutrophils in sites of *L. major* transmission by sand fly bite

As we observed that dermal TRMs and to a lesser extent neutrophils were the main cells that became infected in very early stages after natural transmission, we used intravital microscopy on the upper dermis to reveal the behavior of these cells at sites of *L. major* delivery by infected sand flies. The dorsal skin area of the mouse ear is thin enough for both the epidermis and dermis to be accessed by confocal microscope with long working distance lens. For this, we used LysM-GFP mice to track neutrophils, and Manocept labeling, previously shown to selectively bind mannose receptor (MR) on the surface of dermal TRMs *in situ* [3]. Following infected fly bites, and as previously described [6, 16], we observed recruitment of neutrophils to the site of *L. major* deposition within 1-2 hour post-bite (Fig 2A, Movie S1). All of the parasites appeared immobilized during this time, and they appeared co-localized with neutrophils and/or dermal TRMs, while other parasites had no clear association with either of these cells (Fig 2A). Maximum intensity projection images across *x, y*, and *z* dimensions shows representative magnified images that supports parasite internalization by these respective cells, as well as an example of a parasite that was not co-localized with either of these cell types (Fig 2A, right panels 1, 2 & 3). Another transmission site was imaged, again starting at 1 hr post-bite and continued for 2.5 hr, revealing the swarming of neutrophils in a highly directed manner to the presumed site of vascular wounding (Fig 2B, Movie S2). The number of neutrophils accumulating in the site steadily increased over this time, while their speed and track displacement peaked at 3 hr and decreased at 3.5 hrs post-bite (Fig 2C, D). The total number of parasites in the imaging field did not change over the 2.5 hours of imaging (Fig 2E), and there were a number of parasites that appeared to have been deposited somewhat distal to the bite site that co-localized with dermal TRMs but not with neutrophils (Fig 2B, arrow heads). The dynamic changes in host cell - parasite co-localization indicates that roughly 30% of parasites co-localized with dermal TRMs ahead of infiltrating neutrophils at 1 h post-bite, while at 2-3.5 h the percentage of parasites co-localizing with dermal TRMs and neutrophils increased to roughly 60% and 30%, respectively (Fig 2F).

**Fig 2.**
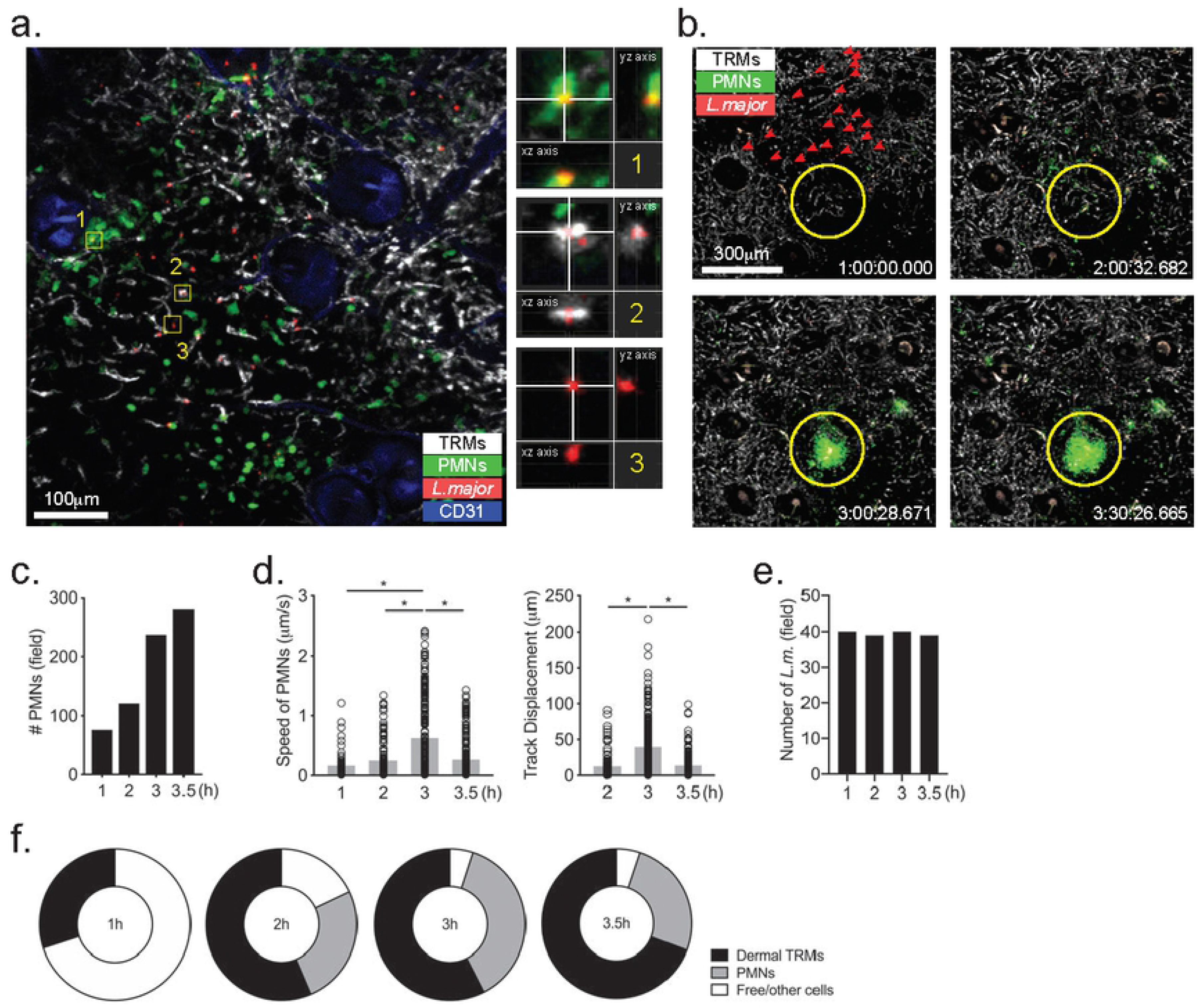
IVM of dermal TRMs and neutrophils in sites of *L. major* transmission by sand fly bite. Sand flies were infected with 5 x 10^6^ RFP^+^ *LmRyn*. After 10-19 days, LysM-GFP mice ears were exposed to infected sand flies. (A) IVM image showing dermal TRMs (white), neutrophils (green), parasites (red) and CD31 (blue) at 1 hr post-transmission. Boxed regions 1, 2, and 3 are enlarged in the panels on the right, showing maximum intensity projection images across *x,y*, and *z* dimensions. (B) IVM time lapse images of another bite site beginning at 1 hr post-bite, showing dermal TRMs (white), neutrophils (green), parasites (red) at 1 hr, 2 hr, 3hr, and 3.5 hr post-bite. (C) Total number of neutrophils in the imaging field at different time points. (D) Speed and track displacement of neutrophils post-sand fly bite. (E) Total number of parasites and (F) percentage of parasites co-localizing with neutrophils or dermal macrophages at different time points. \**P* < 0.05.

Intravital imaging of another transmission site at 3 h revealed a striking example of a high dose transmission in which the parasites were deposited proximal to the area of neutrophil influx (Fig 3A; region 2, Movie S3), and to an area distal from this site (Fig 3A; region 1, Movie S3). While 10-20 parasites in region 2 were co-localized with neutrophils during the 140 minutes of imaging, no co-localization with neutrophils was observed in region 1 (Fig 3B). Imaging of another high dose transmission site at 2 days post-bite again revealed a distinctive pattern of parasite co-localization with neutrophils depending on the area of deposition (Fig 3C & D, Movie 54), with the majority of parasites co-localized with neutrophils in region 1, and few if any co-localized with neutrophils in region 2.

**Fig 3.**
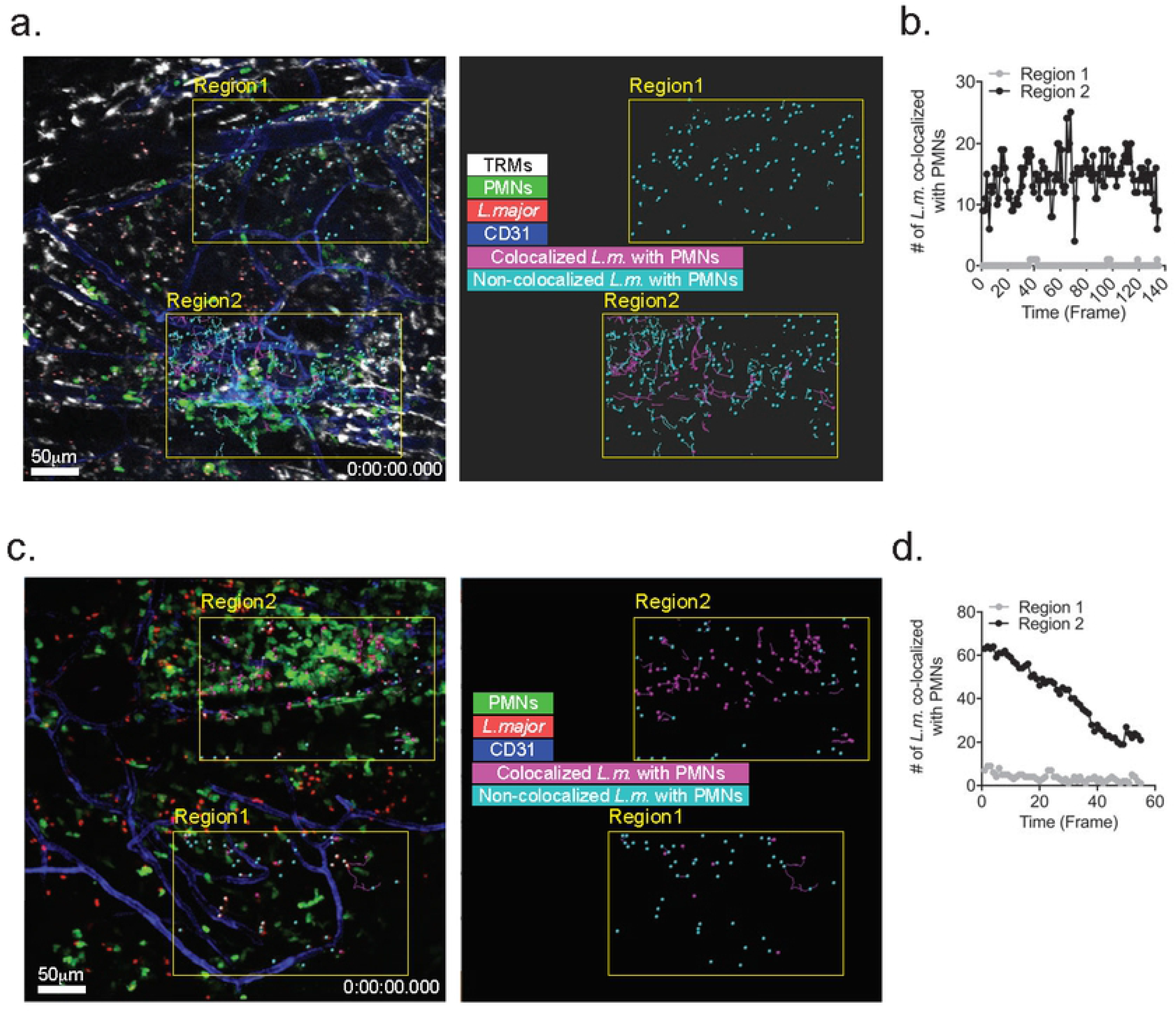
IVM imaging of high dose transmission sites. (A) Confocal image shows transmission site at 3 hr post-bite, with parasites in regions 1 and 2 colorized to indicate their co-localization or not with PMNs. (B) Total number of parasites in regions 1 and 2 co-localized with PMNs during the imaging time span. (C) Confocal image shows transmission site at 2 days post-bite, with parasites in regions 1 and 2 colorized to indicate their co-localization or not with PMNs. (D) Total number of parasites in regions 1 and 2 co-localized with PMNs during the imaging time span.

### Neutrophils transfer *L. major* to dermal macrophages after sand fly transmission

The “Trojan Horse” model of *Leishmania* infection postulates that the uptake of infected, apoptotic neutrophils is a mechanism for the “silent” entry of parasites into macrophages [19, 20]. While the model was originally supported by a series of *in vitro* observations, our prior attempts to capture this process *in situ* by 2P-IVM were unsuccessful [16]. We revisited this model in the context of the interaction of infected neutrophils with dermal TRMs since the later cells had not been properly identified nor adequately labeled in the prior studies. Furthermore, we have shown that dermal TRM efficiently capture apoptotic thymocytes in the skin [3]. By confocal IVM, we could observe infected neutrophils transferring parasites to dermal macrophages (Fig 4A, Movie S5) or undergoing efferocytosis by dermal TRM at 24 hr post-transmission (Fig 4B, Movie S6). The neutrophils that were taken up by the dermal TRMs appeared to be undergoing an apoptotic process, evidenced by membrane blebbing visible in the latter two-time lapse images in Fig 4B. We were also able to visualize dermal TRMs in the process of phagocytosing non-infected, apoptotic PMNs (Fig 4C, Movie S7).

**Fig 4.**
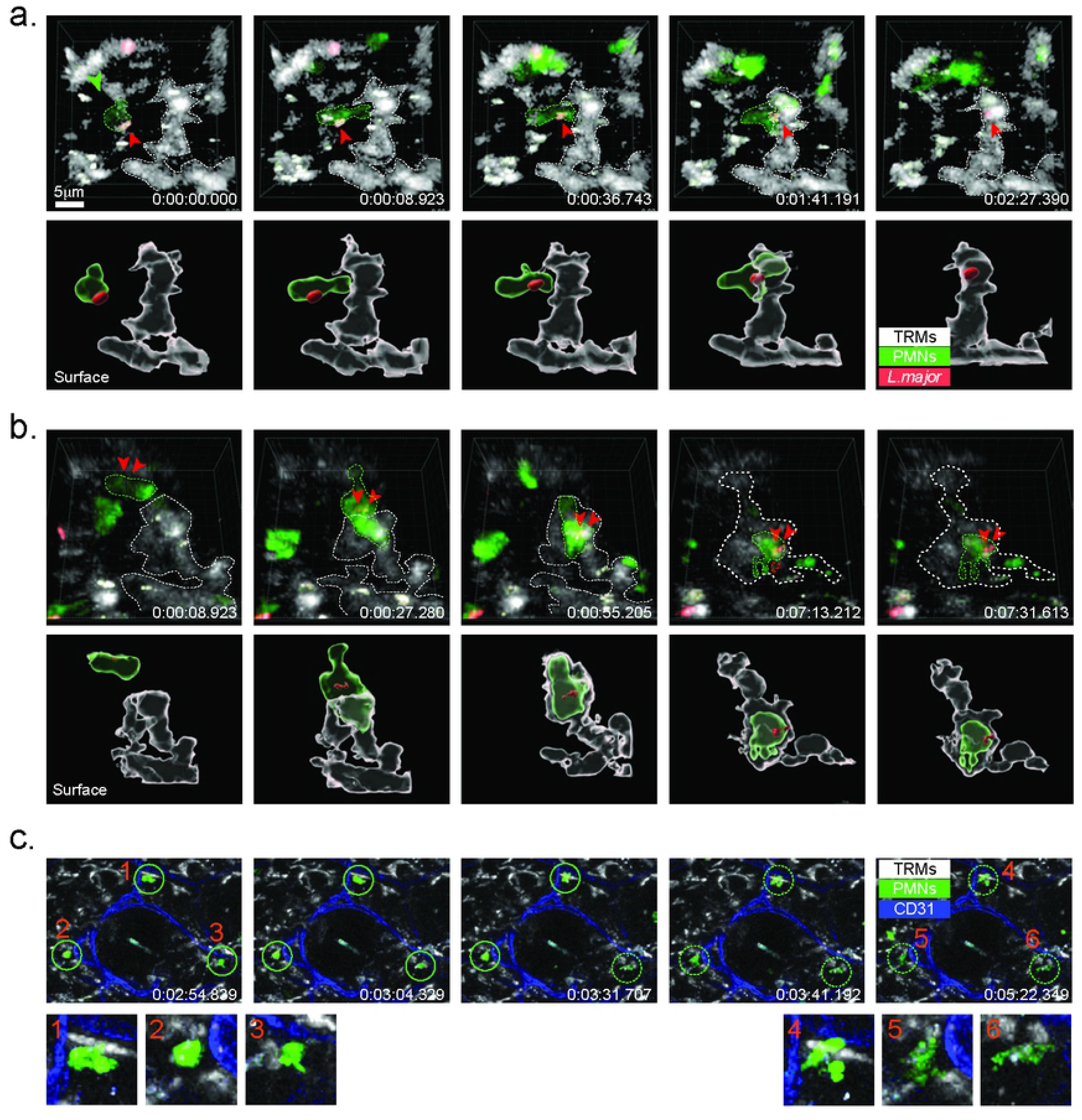
IVM of neutrophils and dermal TRMs interactions. (A) Confocal close-up, time lapse images of infected, LysM-GFP neutrophil interacting with manocept-labeled dermal TRM 24 hrs post-transmission by bite, showing intercellular transfer of RFP *LmRyn*. Bottom row shows 3D surface reconstruction of parasite and host cells. (B) Confocal close-up, time lapse imaging of infected neutrophil interacting with dermal TRM 24 hrs after transmission, showing efferocytosis of infected neutrophil. Bottom row shows 3D surface reconstruction of parasite and host cells. (C). Confocal, time lapse images of non-infected, LysM-GFP neutrophils interacting with manocept-labeled dermal TRMs 24 hrs after transmission. Circled regions are shown in higher magnification below, showing evidence for capture of apoptotic neutrophils by dermal TRMs.

### Dermal TRMs take up apoptotic infected neutrophils *in vivo*

To support the IVM observations regarding the capture of infected neutrophils by dermal TRMs, we infected neutrophils *in vitro* and tracked their uptake by myeloid subsets in the skin following intradermal injection. We have previously shown that following intradermal injection, DCs can take up infected neutrophils and become functionally impaired, but that macrophages are the major population of phagocytes to acquire infection, although at the time we lacked markers to distinguish dermal TRMs from monocyte-derived cells [21]. Infection of neutrophils from bone marrow for 3 hours with several ratios (1:2, 1:5, 1:8) of RFP *LmRyn* yielded a high frequency of infected cells (Fig 5A) that was associated with a higher frequency of apoptotic cells in comparison to uninfected cells in all ratios analyzed (Fig 5B), supporting prior studies regarding the accelerated apoptotic program in neutrophils following their uptake of *L. major* [21]. Neutrophils infected at a 1:8 ratio were loaded with CFSE and injected into the ears of C57Bl/6 mice. The dermal TRMs were the main CFSE^+^ RFP^+^ cells recovered from the ears after 10 min. (Fig 5C and D), indicating that they acquired infection *via* capture of the infected, apoptotic neutrophils. mo-DCs were a minor population of CFSE^+^ RFP^+^ cells, consistent with prior observations [21], while inflammatory monocytes and cDCs appeared to be little involved in this process.

**Fig 5.**
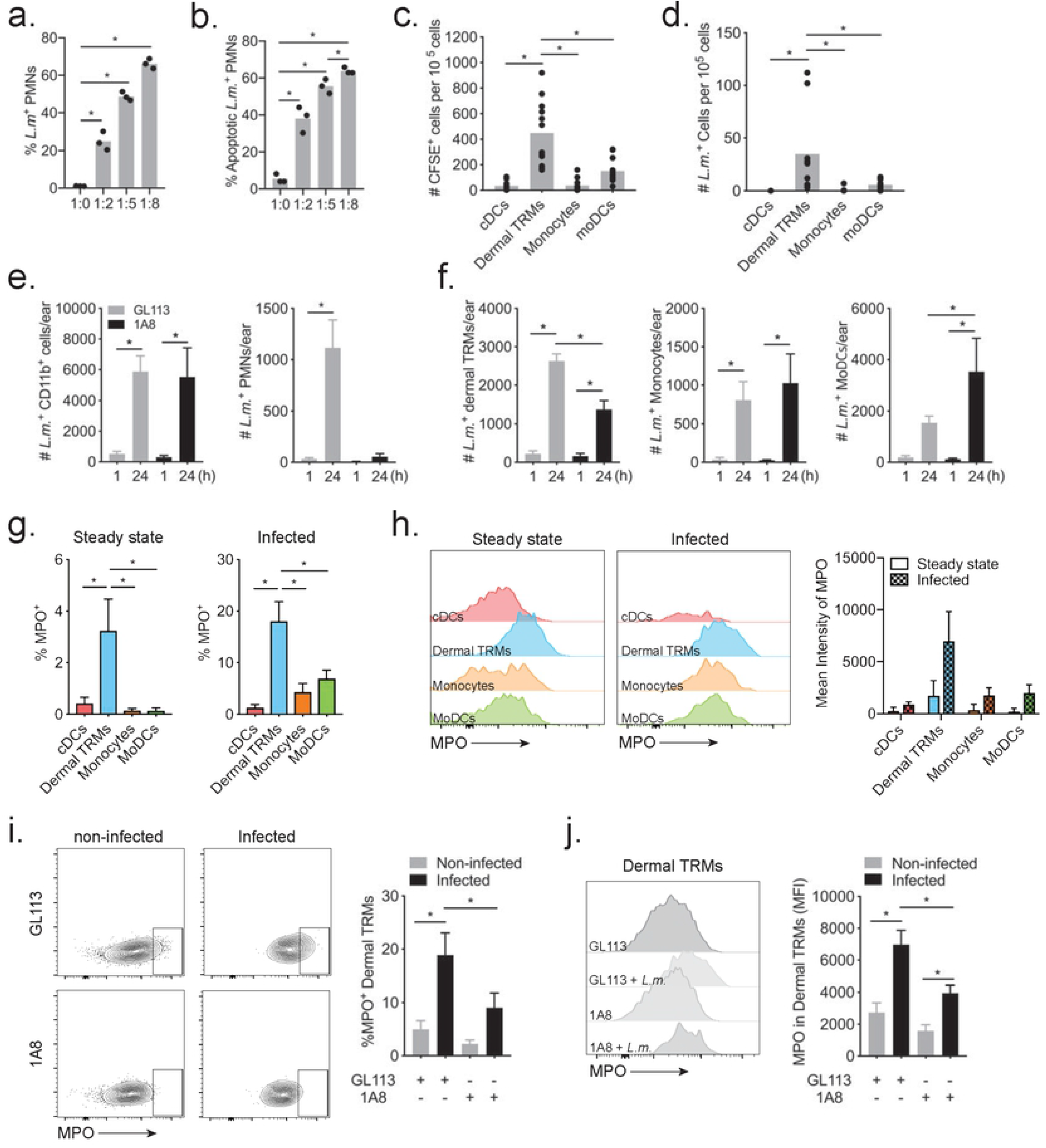
Capture of infected neutrophils by dermal TRMs. A) Frequency of RFP^+^ neutrophils from BM, and (B) frequency of apoptotic neutrophils determined by annexin V staining, following 3 hr incubation with different ratios of RFP+ *LmRyn* metacyclic promastigotes. (C & D) Neutrophils from BM were infected with 1:8 ratio of RFP+ *LmRyn* for 3 hours and then labeled with 1μM CFSE. 2 x 10^5^ neutrophils were injected into ears of C57Bl/6 mice for 10 minutes. Ear dermal cells were harvested and stained for myeloid subsets to determine the number of CFSE+ cells (C) and RFP^+^ cells (D) within each subset. 8-12 ears per group pooled from 2 independent experiments. (E & F) Neutrophil depleted, 1A8 treated or control treated mice were infected with 2 x 10^5^ RFP+ *LmRyn* metacyclic promastigotes in the ear dermis. One hr and 24 hrs post infection, the numbers of infected CD11b+, neutrophils, dermal TRM, inflammatory monocytes and mo-DCs per ear were determined by flow cytometry. Eight ears per group pooled from 2 independent experiments. (G) Frequency of MPO^+^ cells and MPO expression levels with representative histogram plots of MPO staining intensity, in cells from steady state ears and from ears 24 hrs after infection with 2 x 10^5^ RFP+ *LmRyn* metacyclic promastigotes. 12 ears per group pooled from 2 independent experiments. (H) Frequency of MPO^+^ dermal TRMs with representative dot plot, and MPO expression levels in dermal TRMs with representative histogram plots of MPO staining intensity, in cells from 1A8 treated or control treated mice 24 hrs after infection with 2 x 10^5^ RFP^+^ *LmRyn* metacyclic promastigotes. Twelve ears per group pooled from 2 independent experiments. Values shown are means ± SD; \**P* < 0.05.

Because of the low and variable number of parasites deposited by sand fly bite [10], we used intradermal needle inoculation of a relatively high dose (2 x 10^5^) of RFP *LmRyn* metacyclic promastigotes in a series of experiments to quantitatively assess the contribution of parasitized neutrophils to the acquisition of infection by other subsets of myeloid cells in the skin. The pattern of the acute neutrophilic response in these mice is similar to sand fly bite, although more diffuse and comparatively short-lived [6, 16]. Treatment of mice with neutrophil-depleting 1A8 antibody 1 day prior to infection resulted in a 99% and 95% reduction in the number of neutrophils recruited to the ear at 1 hr and 24 hr post-infection (p.i.), respectively (Fig S1A, B). While there was no change in the number of dermal TRMs, the number of inflammatory monocytes and mo-DCs were significantly increased at 24 hr p.i. in the neutrophil depleted mice (Fig S1C, D, and E). Confining the analysis to the infected cells, the total number of Lm^+^CD11b^+^ cells recovered from the site did not change in the neutrophil depleted mice (Fig 5E). As expected, there were no infected neutrophils recovered from these mice. Importantly, the number of infected dermal TRMs was reduced by approximately half at 24 hr p.i. in neutrophil depleted mice compared to the controls (Fig 5F). By contrast, there was a significant, compensatory increase in the number of infected mo-DCs recovered from the neutrophil-depleted mice. These data provide additional evidence that neutrophils contribute to the acquisition of *L. major* infection by dermal TRMs.

To lend further support for a process involving phagocytosis of infected neutrophils, we stained dermal cells for myeloperoxidase (MPO) as a neutrophil-derived marker [21]. In the steady state dermis, approximately 3% of dermal TRM were MPO^+^, while cDCs, inflammatory monocytes and mo-DCs did not present a detectable MPO signal (Fig 5G). Following infection by *L. major*, 17% of infected dermal TRM were MPO^+^, with a significantly elevated expression of MPO compared to their steady state level (Fig 5G). By contrast, only 1% of infected cDCs, 7% of inflammatory monocytes and 4% of mo-DCs were MPO^+^ (Fig. 5G), and their MPO expression levels were significantly less compared to the dermal TRMs (Fig. 5H). Depletion of neutrophils using 1A8 treatment one day prior to *L. major* infection produced a significant reduction in the frequency of infected, dermal TRMs that were MPO^+^ (Fig 5I), and a significant reduction in the MPO expression level on these cells compared to infected, dermal TRMs from control treated mice (Fig 5J).

### Role of receptor tyrosine kinases in infection and alternative activation of dermal TRMs

Accumulating evidence points to a critical role for a subfamily of receptor tyrosine kinases, Tyro3, Axl and Mertk (TAM RTKs), in promoting the resolution of inflammation, including the phagocytic clearance of apoptotic cells [22]. Axl and Mertk, in particular, have been shown to mediate the clearance of apoptotic neutrophils in inflammatory settings [23]. TAM RTKs recognize either directly or indirectly through bridging molecules, the common “eat me” signal of phosphatidylserine (PtdSer), which was shown to be expressed on a high proportion of *in vitro* infected neutrophils, as labeled using annexinV that binds to PtdSer (Fig 5B). Tim-4 is another receptor that can participate in apoptotic cell clearance, including neutrophils, via direct recognition of PtdSer [24]. Analyzing the expression of Axl, Mertk and Tim-4 on dermal TRMs during infection, we found that *L. major* infection upregulated Axl expression, which was otherwise undetectable, but did not significantly modulate the constitutive expression of Tim-4 and Mertk (Fig 6A). We used *Axl^-/-^Mertk^-/-^* mice to explore the involvement of these receptors in the early infection of dermal TRMs following needle inoculation of RFP *LmRyn* in the ear. Infection in the *Axl^-/-^Mertk^-/-^* mice did not significantly modulate the total number of LmRFP^+^CD11b^+^ cells at 1 hr and 24 hr p.i. in comparison to the wild type (WT) mice, nor were the numbers of LmRFP^+^ neutrophils, inflammatory monocytes, or mo-DCs significantly changed (Fig 6B). However, the number of LmRFP^+^ dermal TRMs was significantly decreased by roughly half in the *Axl^-/-^Mertk^-/-^* mice (Fig 6B), which was associated with a small reduction in the total number of dermal TRMs in the injection site at 24 hr p.i., and even in the naive skin (Fig S2).

**Fig 6.**
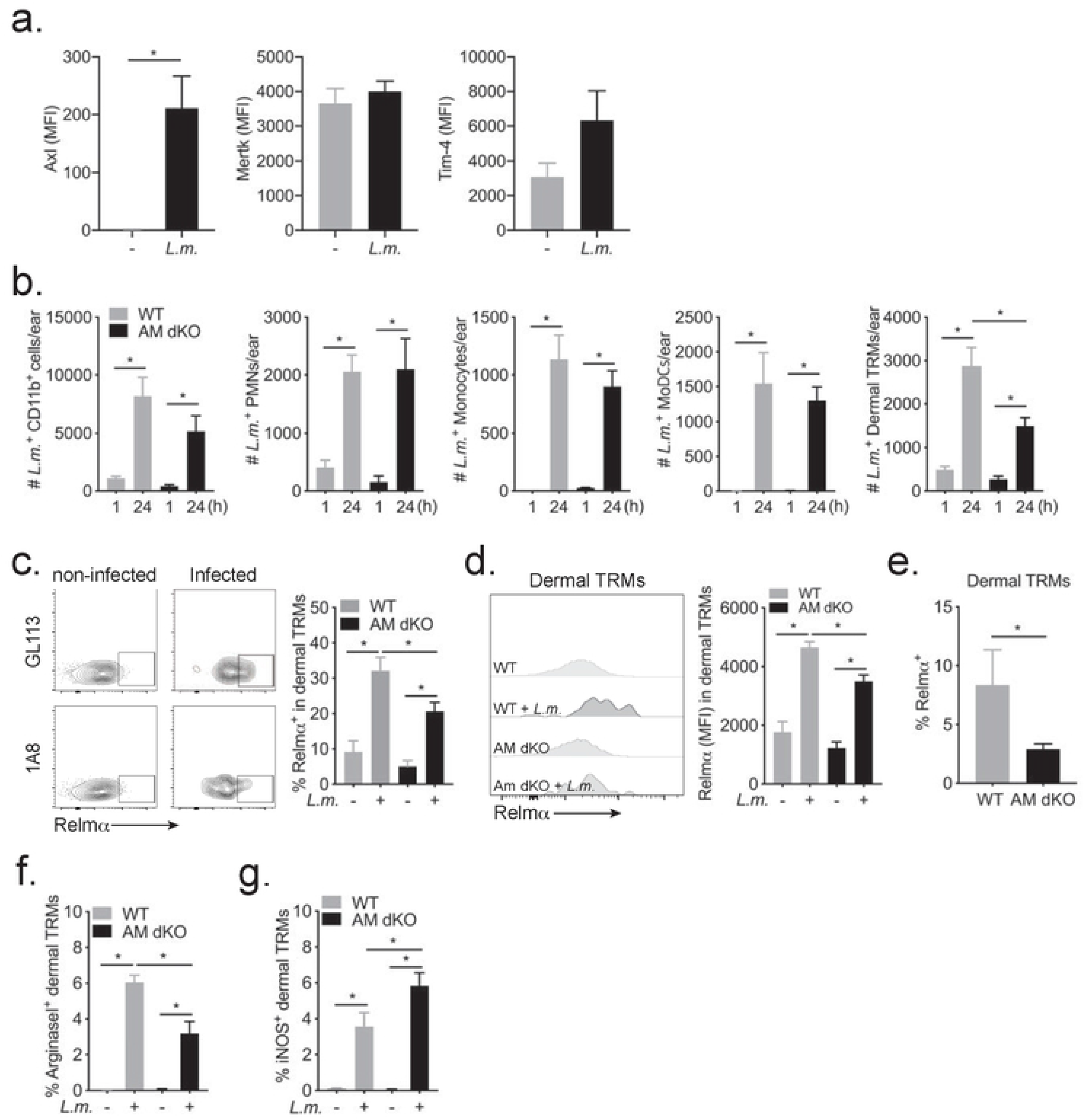
Role of TAM RTKs in infection and M2 polarization of dermal TRM during *L. major* infection. (A) Axl, Tim-4 and MertK expression levels on dermal TRMs from uninfected or infected C57Bl/6 mice 10 days post-infection with 2 x 10^5^ RFP^+^ *LmRyn* metacyclic promastigotes in the ear dermis. 4-6 mice per group. (B) WT and *Axl^-/-^Mertk^-/-^* (AM dKO) mice were infected with 2 x 10^5^ RFP^+^ *LmRyn* metacyclic promastigotes in the ear dermis. One hr and 24 hrs post infection, the numbers of infected CD11b^+^, neutrophils, dermal TRM, inflammatory monocytes and mo-DCs per ear were determined by flow cytometry. 8-16 ears per group pooled from 2 independent experiments. (C) Frequency of Relmα^+^ dermal TRMs and (D) expression level of Relmα with representative histogram plot, in RFP^+^ and RFP^-^ dermal TRMs recovered from the same ear in WT and *Axl^-/-^Mertk^-/-^* mice 48 hr post-infection with 2 x 10^5^ RFP^+^ *LmRyn* metacyclic promastigotes. 12 ears per group pooled from 2 independent experiments. (E) Frequency of Relmα^+^ cells in total dermal TRMs recovered from infected ears in WT and *Axl^-/-^Mertk^-/-^* mice 48 hours post-infection. 12 ears per group pooled from 2 independent experiments. (F) Frequency of Arginase I^+^ and (G) iNOS^+^ cells in RFP^+^ and RFP^-^ dermal TRMs recovered from the same ear in WT and *Axl^-/-^Mertk^-/-^* mice 48 hours post infection; 7-10 ears per group. Values shown are mean frequencies ± SD; \**P* < 0.05.

The anti-inflammatory, pro-resolving functions of macrophages involves the upregulation of specific gene expression programs, of which *relmα* and *arg1* are considered signature components [25]. Comparing the expression of Relmα on LmRFP^-^ and LmRFP^+^ dermal TRMs recovered from the same site at 48 hr p.i. in WT mice, 9% and 32% of these cells, respectively, stained positive for Relmα (Fig 6C). Relmα expression levels were also significantly higher in the infected vs. uninfected dermal TRMs (Fig 6D). In the *Axl^-/-^Mertk^-/-^* mice, the frequency of Relmα^+^ cells and the Relmα expression levels were also significantly higher in the infected compared to uninfected dermal TRMs recovered from the same site (Fig 6C and D). These parameters, however, were in each case significantly reduced compared to WT mice (Fig 6C and D), resulting in a significant decrease in the frequency of Relmα^+^ cells in the total dermal TRM population in the *Axl^-/-^Mertk^-/-^* mice (Fig 6E). A similar consequence of the *Axl/Mertk* deficiency was observed with regard to the expression of Arg1 on dermal TRMs from infected mice, which was confined to infected cells, and reduced in frequency on infected cells from the *AxP^-/-^Mertk^-/-^* mice (Fig 6F). Arg1 expression was also upregulated on infected inflammatory monocytes and mo-DCs, which did not, however, appear to be dependent on Axl/Mertk (Fig S3). The reduced frequency of Arg1^+^ cells in the population of infected, dermal TRMs from *Axl^-/-^Mertk^-/-^* mice was associated with a significantly increased frequency of iNOS^+^ cells in this population (Fig 6G). Again, no effect of the Axl/Mertk deficiency was observed on the expression of iNOS in the populations of infected inflammatory monocytes and mo-DCs (Fig S3). Taken together, these data demonstrate that TAM RTKs contribute to the early infection and M2-like activation program of dermal TRMs following *L. major* infection in the skin.

### Role of dermal TRMs and TAM RTKs in *L. major* infection outcome *in vivo*

We tested the early infection outcome in mice transiently depleted of dermal TRMs following treatment with M279, an antibody to colony stimulating factor-1 receptor (CSF-1R). As previously shown [3], the dermal TRMs were confirmed to be depleted without significant off-target effects on other lymphoid or myeloid populations in the skin (Fig 7A). Because it was not possible to maintain the depletion of dermal TRMs for an extended period of time, we analyzed early parasite burdens in mice infected with high dose needle challenge. Dermal TRMs-depleted mice had a significant 4-fold lower parasite burden at d3 and 5-fold lower burden at d9 p.i. (Fig 7B), confirming their contribution to the early establishment of infection in this challenge model.

**Fig 7.**
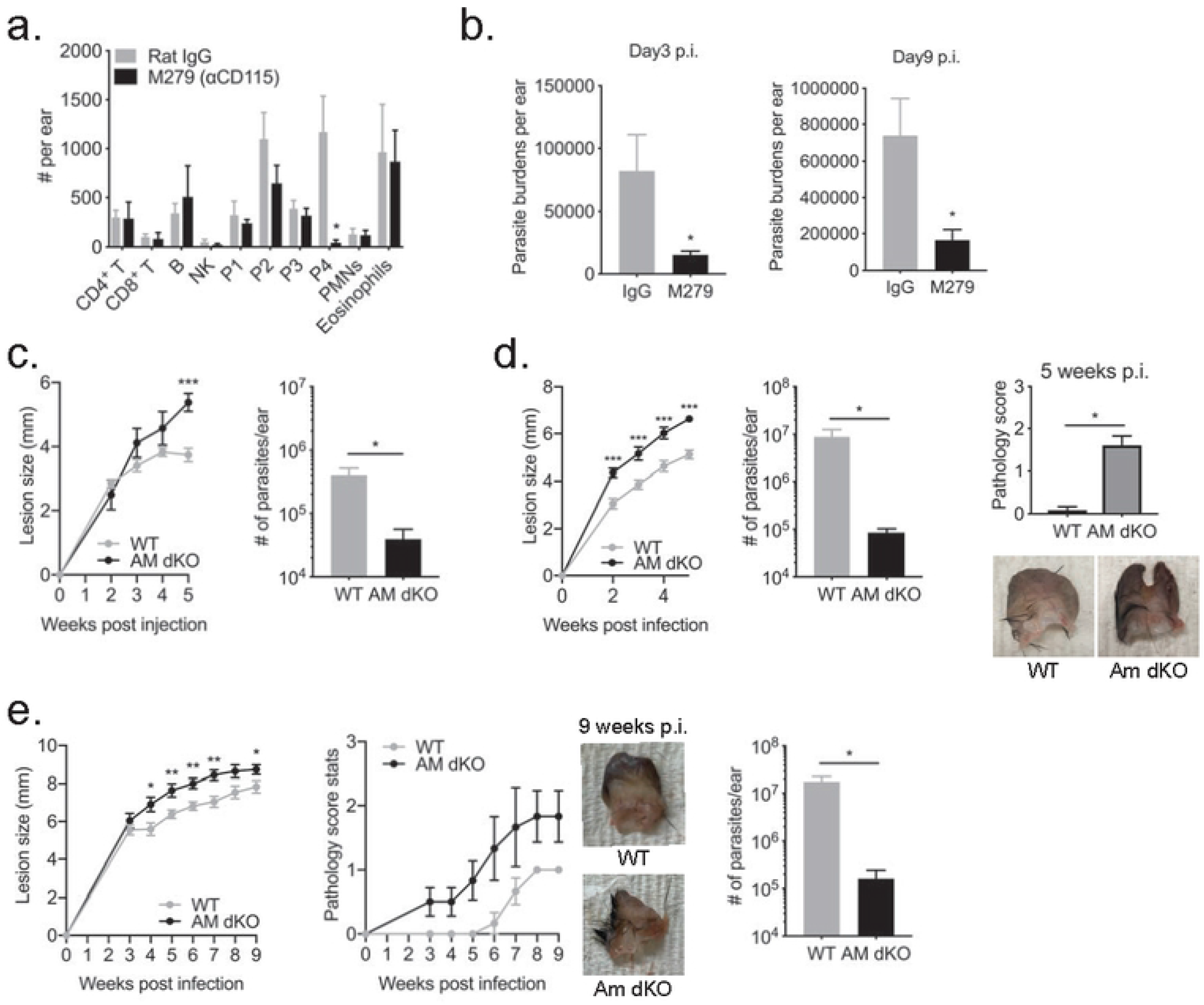
Dermal TRMs and TAM RTKs contribute to infection outcome *in vivo*. (A) The total number of lymphoid and myeloid subsets (P1-inflammatory monocytes, P2-MoDC, P3-cDC, P4-dermal TRM) recovered from the ear after treatment with M279 three times a week for 3 wk; 4 ears/group. (B) Parasite burdens at 3 and 9 days p.i. in the ear dermis of C57Bl/6 mice treated with either M279 or control IgG three times a week for 3 weeks and then infected with 2 x 10^5^ *LmRyn* metacyclic promastigotes; 6 ears/group at each time point. (C) WT and *Axl^-/-^Mertk^-/-^* mice (AM dKO) were infected with 2 x 10^5^ *LmRyn* metacyclic promastigotes in the ear dermis. Lesions size was measured weekly during 5 weeks infection and lesion parasite burden determined at 5 weeks. 12 ears per group. (D) Neutrophils from BM were infected at a 1:8 ratio with *LmRyn* metacyclic promastigotes for 3 hours and injected into ears of WT and *Axl^-/-^Mertk^-/-^* mice. Lesion size was measured weekly, and both pathology score and lesion parasite burden determined at 5 weeks p.i.; 12 ears per group. (E) *Phlebotomus duboscqi* sand flies were infected with 5 x 10^6^ RFP^+^ *LmRyn* procyclic promastigotes. After 10-19 days, ears of WT and *Axl^-/-^Mertk^-/-^* mice were exposed to infected sand flies. Lesions size and pathology score were measured weekly during 9 weeks infection and lesion parasite burden determined at 9 weeks; 8 ears per group. Values shown in A-E are means ± SD; \**P* < 0.05.

Finally, WT and *Axl^-/-^Mertk^-/-^* mice were infected with RFP *LmRyn* metacyclic promastigotes by needle in the ear dermis. The size of the nodular lesions in the *Axl^-/-^Mertk^-/-^* mice were larger than the WT mice at 5 weeks p.i. (Fig 7A). Interestingly, the differences in lesion severity did not correlate with parasite burden, which was approximately 10 times lower in the ears from the *Axl^-/-^Mertk^-/-^* mice at 5 weeks p.i. When the infections were initiated using parasitized, apoptotic neutrophils, we observed larger lesions in the *Axl^-/-^Mertk^-/-^* mice starting at 2 weeks p.i. and more severe lesion pathology, as reflected by the degree of ulceration and tissue erosion (Fig 7D). Despite the severity of their lesions, parasite burden was approximately 100 fold lower in the *Axl^-/-^Mertk^-/-^* mice. Lastly, we infected sand flies for natural transmission of RFP *LmRyn* to ears of WT and *Axl^-/-^Mertk^-/-^* mice. As observed in the needle inoculations, the *Axl^-/-^Mertk^-/-^* mice showed larger lesions throughout the course of infection, and their pathology scores were substantially exacerbated (Fig 7E). Again, the parasitic load in the lesion was roughly 100 times lower in the *Axl^-/-^Mertk^-/-^* mice (Fig 7E). These data demonstrate the importance of TAM RTKs in both the establishment of *L. major* infection after natural transmission and their role in tissue repair during infection.

## Discussion

The current studies have used flow cytometry of ear dermal cells and confocal IVM of the upper dermis to reveal the early events following the delivery of *L. major* into the skin by the bite of an infected natural vector, *P. duboscqi*. To acquire a bloodmeal from the vertebrate host, female sand flies lacerate superficial capillaries in the dermis to create a pool of blood upon which they feed. In addition, sand flies salivate into the tissue to inhibit hemostasis, and can egest other molecules that have immunomodulatory properties [14]. In infected flies, the egestion of parasites into the skin is thought to occur as a consequence of heavy infections in the foregut or anterior midgut that interfere with the normal directional flow of the meal through the food canal [26], or that can form a plug comprised of promastigotes embedded in a gel-like matrix, resulting in regurgitation of parasites during attempts by the fly to dislodge the plug [27]. These feeding conditions can produce considerable variability in the number of transmitted promastigotes and in the types of phagocytes to which transmitted parasites are exposed at the outset of infection. By flow cytometry analysis, dermis TRMs were on average the predominant infected cell type at 1 hr and 24 hr, although there was considerable variability in these proportions depending on the site. Neutrophils, monocytes, and mo-DCs were also among the infected cells during these early time points, with neutrophils, like dermal TRMs, representing an especially variable proportion of the infected cells. A better understanding of the nature of this variability was afforded by the intravital imaging of different transmission sites, which revealed deposition of parasites that in some cases were distal to the presumed area of vascular damage, with little or no co-localization with neutrophils. By contrast, parasites deposited in close proximity to the actual bite site, demarcated by the highly localized swarming of neutrophils, were frequently co-localized with and taken up by neutrophils, similar to the original observations made using 2P-IVM [16]. It is not known if there are differences in the feeding behavior or infection status of the flies that results in their delivery of parasites at some distance from the hemorrhagic pool from which they feed. The source of these parasites may be flies that have probed but abandoned their feeding attempts in that particular site in favor of another attempt in a proximal site.

The role of dermis TRMs in the early events following transmission by bite has not been previously investigated, and follows on recent studies that have characterized their embryonic origin, radio-resistance, and perivascular distribution in the steady state murine dermis [3, 17]. Furthermore, dermal TRMs are highly phagocytic and retain M2-like transcriptional profiles even under strong pro-inflammatory conditions, consistent with their role in promoting tissue homeostasis and repair [3, 17]. Thus, they appear to be pre-positioned and pre-conditioned to serve as host cells for the initial uptake and survival of *Leishmania* in the inoculation site. The especially high proportionate representation of the dermal TRMs among the infected cells at 1 and 24 hrs in some of the transmitted ears may reflect the fate of parasites deposited distal to the hemorrhagic site, discussed above, for which competing encounters with infiltrating neutrophils and monocytes may not occur. Starting at 5 days post-transmission, and especially by 12 days, the infections transitioned into mainly inflammatory cells, although there was considerable variability in the myeloid subsets infected, emphasizing the heterogeneity of natural transmission sites even at 1-2 weeks p.i. At 5 days, a few of the ears showed a relatively high proportionate number of infected eosinophils, extending their influx to natural transmission sites, which was shown to be mediated by CCL24 produced by dermal TRM following infection by needle [28]. Neutrophils and mo-DCs each showed an especially high variability in their proportionate representation of infected cells at these time points. The variability in the numbers of infected inflammatory monocytes and mo-DCs may reflect differences in the onset of an adaptive Th1 response, shown to be required for monocyte recruitment following infection by needle [29].

At early time points, in addition to their direct uptake of parasites, some of the dermal TRMs may have acquired their infections via transfer from or uptake of parasitized neutrophils, i.e. the “Trojan Horse” model, a process that we believe was captured in real time by IVM (Fig 4, Movies S5 and S6). The ability of the dermal TRMs to efferocytose infected neutrophils in the transmission site is consistent with their unique ability to efficiently capture apoptotic cells [3]. When neutrophils from BM were infected with RFP *LmRyn* and CFSE labeled *in vitro*, the majority rapidly acquired an apoptotic marker, and were taken up primarily by dermal TRMs within 10 min following their injection into the ear dermis (Fig 5A-D). Our prior study also identified macrophages as the predominant infected population following injection of infected neutrophils, although at the time we were unable to phenotype the cells as dermal TRMs [21]. In addition, while the cells in the prior study were positive for the parasite they were negative for the neutrophil marker. This is likely explained by the longer interval between injection and recovery of the ear dermal cells for analysis (4 hrs) that may have resulted in phagosomal degradation of the neutrophil GFP signal following engulfment, or by the direct transfer of parasite from infected neutrophils to dermal TRMs, as revealed in the IVM.

Additional evidence supporting a specialized role for dermal TRMs in capture of infected neutrophils *in vivo* is the effect of neutrophil depletion just prior to infection that resulted in an approximate 50% reduction in the number of infected dermal TRMs at 24 hr, while the numbers of infected monocytes and mo-DCs were increased (Fig 5E). Finally, a large number of dermal TRMs became positive for the neutrophil marker MPO during infection, while this number was reduced in neutrophil depleted mice (Fig 5F & G). The caveat with the neutrophil depletion experiments is that they were performed in mice with relatively high dose infections delivered by needle to contend with the high variability in both the numbers of fly transmitted parasites and in their co-localization with neutrophils. By contrast, needle inoculations of *Leishmania* in the ear dermis have consistently shown that neutrophils predominate as the initial infected cell in the site [6, 16, 21, 30, 31]. While the current studies emphasize that sand fly transmissions only occasionally conform to this outcome, the artificial infections nonetheless reinforce the evidence from the limited IVM observations that for the neutrophils that do become infected following fly transmission, their engulfment by and transfer of parasites to dermal TRMs is a likely scenario.

The efficient clearance of abnormal or dying cells, including neutrophils, by macrophages and DCs is a crucial homeostatic process, and is largely dependent on the exposure of PtdSer on the outer leaflet of the membrane, an evolutionary conserved “eat-me” signal for apoptotic cells. PtdSer is recognized by a number of different receptors, among which the TAM RTKs, together with their cognate agonists, play an especially important role in the resolution of inflammation [22, 32]. In mice, Mertk is expressed on most mature tissue macrophages, whereas Axl is more tissue specific, although it can be upregulated by inflammatory stimuli [33]. Our findings confirm the expression of Mertk on dermal TRMs from naïve mice, and show that Axl is expressed only on dermal TRMs recovered from infected mice (Fig 6a). *L. major* infection (by needle) of *Axl^-/-^Mertk^-/-^* mice led to a significant reduction in the number of infected dermal TRMs at 24 hr (Fig 6B), similar to the effect of neutrophil depletion, and consistent with the involvement of Axl and Mertk in the efferocytosis of infected neutrophils. Apoptotic cell clearance by PtdSer-dependent TAM RTKs is linked to the induction of an anti-inflammatory program, and in conjunction with IL-4/IL-13 has been shown to maintain the reparative program of tissue resident macrophages [34]. Relmα and Arginase1 in particular, contribute to the initiation of pro-fibrotic pathways and other tissue-remodeling functions [35, 36]. The expression levels and frequency of cells positive for each of these markers were significantly increased on infected compared to non-infected dermal TRMs recovered from the same site. While this was also true of cells recovered from the *Axl^-/-^Mertk^-/-^* mice, compared to wild type mice there were lower frequencies of Relmα^+^ or Arginase1^+^ cells and a greater frequency of iNOS^+^ cells within the population of infected, dermal TRMs. These differences were associated with clear effects on infection outcomes in the *Axl^-/-^Mertk^-/-^* mice, whether initiated by infected neutrophils or metacyclic promastigotes delivered intradermally by needle, or most critically by infected sand flies. In each case, the *Axl^-/-^Mertk^-/-^* mice showed more severe pathology despite substantially fewer numbers of parasites in the lesion. Thus, the dermal TRMs, dependent at least in part on their expression of Axl and Mertk, appear to provide an early replicative niche for the parasite while at the same time contributing to the resolution of inflammation and tissue repair. In other infection models, a role for TAM RTKs in promoting critical macrophage efferocytotic functions have been reported. Axl-deficient mice challenged with influenza virus showed enhanced morbidity, associated with reduced efferocytotic capacity of aveolar macrophages and accumulation of dead cells in the lung [37], and Axl induction during viral infection helped to maintain the ability of human macrophages to efferocytose apoptotic cells [38]. It is important to keep in mind that the phenotype observed in the *L. major* infected *Axl^-/-^Mertk^-/-^* mice may not have been directly related to deficits in apoptotic cell clearance, since other consequences of TAM receptor signaling, such as negative regulation of dendritic cell function, have been described [39].

In summary, our analyses of sand fly transmission sites of *L. major* in the mouse skin revealed that dermis TRMs were the predominant phagocytes to take up the parasite within the first 24 hr post-bite. The early involvement of neutrophils varied depending on the proximity of deposited parasites to the site of vascular damage around which the neutrophils coalesced. Some of the dermal TRMs could be visualized acquiring their infections via transfer from or efferocytosis of parasitized neutrophils, providing direct evidence for the “Trojan Horse” model. The involvement of TAM RTKs in these interactions and in sustaining the pro-resolving functions of dermal TRMs was supported by reduced parasite burdens but more severe pathology in *Axl^-/-^Mertk^-/-^* mice. The heterogeneity of sand fly transmission sites with respect to dose and the early cellular interactions described here likely contribute to the wide range of infection outcomes that are associated with natural transmission of *L. major* observed in mouse models [16, 40, 41], and possibly humans.

## Methods

### Mice

C57Bl/6, C57Bl/6 LysM-GFP and BALB/c mice were obtained from Taconic Laboratories and *Axl^-/-^Mertk^-/-^* mice on a C57Bl/6 background were kindly provided by Dr. Carla Rothlin (Yale University School of Medicine). All mice were kept under pathogen-free conditions in the NIAID animal care facility with sterilized water, shavings and commercial rations. All mice used in this work were female and the study protocol was approved by the NIAID Animal Care and Use Committee (no. LPD 68E). Use of animals in this research was strictly monitored for accordance with the Animal Welfare Act, the Public Health Service Policy, the U.S. Government Principles for the Utilization and Care of Vertebrate Animals Used in Testing, Research, and Training, as well as the National Institutes of Health Guide for the Care and Use of Laboratory Animals.

### Parasites

*Leishmania major* Ryan strain, originating in Iraq and stably transfected with a red fluorescent protein (RFP *LmRyn*), has been described previously [18]. The promastigotes were grown at 26°C in 199 media supplemented with 20% fetal bovine serum (FBS), 100 units/mL penicillin, 100 μg/mL streptomycin, 2 mM L-glutamine, 40 mM Hepes, 0.1 mM adenine (50 mM Hepes), 5 mg/mL hemin (50% triethanolamine), and 1 mg/mL 6-biotin (M199/S). RFP *LmRyn* was grown in the presence of 50 μg/ml Geneticin (G418) (Sigma). For sand fly infections, parasites were used in log phase (1-2 day culture). For mouse infection, infective metacyclic promastigotes were isolated from stationary phase cultures (5 to 7 days) by Ficoll^®^ density gradient centrifugation as previously described [42], and injected in the ear dermis in a volume of 10 μL.

### Sand fly infections and transmission by bite

Two-to-four day old *Phlebotomus duboscqi* females were obtained from a colony initiated from field specimens collected in Mali. Flies were infected by artificial feeding through a chick skin membrane on heparinized mouse blood seeded with 4 x 10^6^ / ml RFP *LmRyn* promastigotes, as previously described [18]. The separated mouse plasma was heat inactivated at 56°C for 1 hr. prior to adding back to the packed red blood cells. Blood engorged flies were separated and maintained at 26°C and 75% humidity and were provided 30% sucrose *ad libitum. Leishmania* infections were allowed to mature for 9-11 days within the sand fly midgut. One day before transmission the sucrose diet was removed. On the day of transmission, 10 flies were transferred to small plastic vials covered at one end with a 0.25-mm nylon mesh. Mice were anesthetized by intraperitoneal injection of 30 ul of ketamine/xylazine (100 mg/ml). Specially designed clamps were used to bring the mesh end of each vial flat against the ear, allowing flies to feed on exposed skin for a period of 2 hours in the dark. In each transmission experiment, some mouse ears were also exposed to the bites of uninfected flies.

### Determination of lesion size and parasite load

Following needle inoculation of metacyclic promastigotes or exposure to infected sand flies, ear lesion diameters were measured (in mm) weekly for up to 9 weeks, and pathology was scored as previously described [43], using the following scale: 0 = no ulcer, 1 = ulcer, 2 = ear half eroded, 3 = ear completely eroded. For quantification of parasite load in the infected ear, parasite titrations were performed on tissue homogenates as previously described [43]. The number of viable parasites in each ear was determined from the highest dilution at which promastigotes could be grown out after 7–10 days of incubation at 26°C.

### Neutrophils isolation, *in* vitro infection, adoptive transfer, and *in vivo* depletion of neutrophils and dermal TRMs

Neutrophils from bone marrow were purified from C57Bl/6 mice by negative selection using neutrophil isolation kit (Miltenyi Biotec). Purified neutrophils were cultured at 1.0-5.0 x 10^7^ total cells and were infected or not at different ratios with RFP *LmRyn* metacyclic promastigotes, serum opsonized by prior incubation for 30 min. in 5% normal mouse serum. After 3 hr incubation at 35°C, 5% CO_2_, infection levels (RFP) and expression of apoptotic marker (Apoptosis Assay, Thermo Fisher) were quantified by FACS. For mice infection, neutrophils from bone marrow were infected with RFP *LmRyn* metacyclic promastigotes 1:8 ratio for 3 hours, and 2.0 x 10^5^ neutrophils were injected in the ears of C57Bl/6 and *Axl^-/-^Mertk^-/-^* mice. Neutrophils were depleted using a single i.p. injection of 1 mg of 1A8 (anti-Ly6G, BioXCell), or GL113 (control IgG, BioXCell). One day after treatment, ears were injected with 2 x 10^5^ RFP *LmRyn* metacyclic promastigotes in the dermis by i.d. injection in a volume of 10 μL. M279 (Amgen) is a rat IgG mAb, which blocks ligand binding to the CSF-1R. Mice were treated with 200 μg M279 or rat IgG (Sigma) intraperitoneally 3 times a week for 9 weeks. The efficiency and specificity of the depletions were evaluated in dermal cell preparations by FACS.

### Processing of ears, skin cell immunolabeling and flow cytometry analysis

Uninfected and infected ear tissue was harvested as previously described [44]. Briefly, the two sheets of the ears were separated, placed in DMEM containing 0.2 mg/mL Liberase Cl purified enzyme blend (Roche Diagnostics Corp.), and incubated for 1.5 hr at 37°C. The digested ears were processed in tissue homogenizers (Medimachine; BD Bioscience) and filtered through a 70μm pore cell strainer (BD Biosciences). Single-cells suspensions were stained with 1μM Live/Dead Aqua Dead Cell Stain Kit (ThermoFisher) for 30 min at 4°C, followed by incubation with 50 ng/mL anti-Fc-γ III/II (CD16/32) receptor Ab (93, BioLegend) in 100 μl of FACS Buffer (1% FBS and 1 mM EDTA in PBS). The fluorochrome-conjugated antibodies (20 ng/mL) were added for 30 minutes at 4°C. The following antibodies were used for surface staining: PE-Cy7-anti-mouse CD11b (M1/70, BioLegend); FITC-anti-mouse Ly6G (1A8, BioLegend); APC-Cy7-anti-mouse Ly6C (HK1.4, BioLegend); Brilliant Violet 421-anti-mouse SiglecF (E50-L440, BD Bioscience); PerCP-Cy5.5-anti-mouse CD64 (X54-5/7.1, BioLegend); APC-anti-mouse CD206 (C068C2, BioLegend). For intracellular staining, PE-anti-mouse MPO (8F4, Hycult); PE-anti-mouse iNOS (CXNFT, Invitrogen); PE-anti-mouse Arginase I (Met1-Lys322, R&D Systems), PE-anti-mouse Relmα (D58RELM, Invitrogen) were used. For intracellular staining, the cells were stained for their surface markers, then fixed and permeabilized using BD Cytofix/Cytoperm (BD Biosciences) and finally stained for detection of intracellular targets by incubation for 30 min on ice. The isotype controls used were PE-IgG1 (RTK2071, BioLegend); PE-IgG2α (RTK2758, BioLegend); and IgG2β (eB149/19H5, eBioscience). The cells were washed 2 times with FACS Buffer and the data collected using FacsDIVA software (BD Bioscience) and a FacsCANTO II flow cytometer (BD Biosciences), with acquisition of at least 50,000 events. The number of cells were estimated using AccuCheck Counting Beads (Life Technologies). The data were analyzed using FlowJo, LLC software (BD Bioscience).

### Intravital microscopy

Intravital microscopy of the sand fly transmission sites in C57Bl/6 LYS-eGFP mice was performed as previously described [28], using confocal microscopy *in vivo*. The mice were intravenously injected with 20μg of eFluor450 anti-mouse CD31(390, Invitrogen), and with 25μg Manocept-Cy5 or -Alexa488 to visualize dermal TRMs, immediately prior to imaging. Non-invasive intravital imaging of mouse ear was performed using Leica DIVE (Deep In Vivo Explorer) inverted confocal microscope (Leica Microsystems) with full range of visible lasers (Spectra Physics). Additionally, the microscope was equipped with ultra-sensitive DIVE detector; Lx25.0 water-immersion objective with 0.95 NA (Leica Microsystems) and 2 mm working distance; a motorized stage; and Environmental Chamber (NIH Division of Scientific Equipment and Instrumentation Services) to maintain 37° C. Anesthesia was induced with 2 % Isoflurane (Baxter) and maintained at 1.5 % during imaging. Mouse ears were immersed in carbomer-based solution to prevent dehydration, and fixed flat using surgical tape on the cover-glass bottom stage for imaging. A temperature sensor was positioned on the stage near the animal. Mai Tai was tuned to 880 nm excitation; InSigth was tuned to 1150 nm. Diode laser was used for 405 nm excitation; Argon laser for 488 nm excitation; DPSS laser for 561 nm excitation; and HeNe laser for 633 nm excitation wavelengths. All lasers were tuned to minimal power (between 0.5 and 3 %). For time-lapse imaging, small tiled images of 2×2 fields were recorded over time. Z stacks consisting of 6-8 single planes (3-5 μm each over a total tissue depth of 50-70 μm) were acquired every 45 seconds for a total observation time between 1 to 6 hours for 4D reconstruction, surface modeling and tracking with the Imaris software (Imaris version 9.2.1, Bitplane AG, Zurich, Switzerland). Cells were segmented as 3D surface model and tracked using Imaris autoregressive tracking algorithm. Cell tracks are shown as lines. Colocalization between cells and parasites was calculated using “Kiss and Run Analysis” extension for Imaris. Distance Transformation outside of the target surface object method was used to determine closest surface to surface distance of the tracked objects.

### Statistical analysis

The differences in values obtained for two different groups were determined using non-parametric Mann-Whitney test. For comparisons of multiple groups, one-way analysis of variance (ANOVA) followed by Dunn’s post-test was used. Analyses were performed using Prism 7.0 software (GraphPad).

## Acknowledgments

We thank Dr. Carla Rothlin, Yale University School of Medicine, for the provision of the *Axl^-/-^Mertk^-/-^* mice, and Dr. Nathan Peters, University of Calgary, for helpful comments and discussions.

## Author contributions

MMC, SHL, and DS conceived and designed the experiments. MMC, SHL, OK, and KG performed the experiments. MMC, SHL, and DS wrote the paper.

## Supporting information

**S1 Fig. Depletion of neutrophils modulates myeloid populations in the skin after *L. major* infection**. 1A8 treated or control treated C57Bl/6 mice were infected 24 hr later with 2 x 10^5^ RFP^+^ *LmRyn* metacyclic promastigotes in the ear dermis. One hr and 24 hrs p.i., the numbers of CD11b^+^ subsets were determined by flow cytometry. (A) Representative dot plots of ear dermal cells, (B) neutrophils, (C) dermal TRM, (D) inflammatory monocytes and (E) mo-DCs per ear were determined by flow cytometry. Values shown are mean number of cells per ear ± SD, 8-12 ears per group pooled from 2 independent experiments; \**P* < 0.05.

**S2 Fig. Receptor tyrosine kinases modulate myeloid populations in the skin after *L. major* infection.** (A-D) WT and *Axl^-/-^Mertk^-/-^* (AM dKO) mice were infected with 2 x 10^5^ RFP+ *LmRyn* metacyclic promastigotes in the ear dermis. One hr and 24 hrs post infection, the numbers of dermal TRM, neutrophils, inflammatory monocytes and mo-DCs per ear were determined by flow cytometry. Values shown are mean number of cells per ear ± SD, 8-12 ears per group pooled from 2 independent experiments; \**P* < 0.05.

**S3 Fig. Arginase 1 and iNOS are not modulated by Axl and Mertk receptor tyrosine kinases in inflammatory monocytes and mo-DCs.** WT and *Axl^-/-^Mertk^-/-^* (AM dKO) mice were infected with 2 x 10^5^ RFP^+^ *LmRyn* metacyclic promastigotes in the ear dermis. Forty eight hours after infection, frequencies of ArginaseI^+^ and iNOS^+^ cells in inflammatory monocytes (A, C) and mo-DCs (B, D) were determined by flow cytometry. Values shown are mean number of cells per ear ± SD, 7-12 ears per group pooled from 2 independent experiments; \**P* < 0.05.

**Movie S1: IVM showing recruitment of neutrophils to the site of *L. major* transmission, 1-2 hr post-bite.**

**Movie S2: IVM showing swarming of neutrophils to the presumed site of vascular wounding, 1-2.5 hr post-bite.**

**Movie S3: IVM of high dose transmission site, 3 hr post-bite.**

**Movie S4: IVM of high dose transmission site, 2 days post-bite.**

**Movie S5: IVM of infected neutrophils transferring parasites to dermal macrophages, 24 hr post-bite.**

**Movie S6: IVM of infected neutrophils undergoing efferocytosis by dermal TRM, 24 hr post-bite.**

**Movie S7: IVM of dermal TRMs in the process of phagocytosing non-infected PMNs, 24 hr post-bite.**

